# Methyl-CpG binding domain protein 2 plays a causal role in breast cancer growth and metastasis

**DOI:** 10.1101/2020.10.06.328989

**Authors:** Niaz Mahmood, Ani Arakelian, Moshe Szyf, Shafaat A. Rabbani

**Affiliations:** Department of Medicine, McGill University, Montréal, QC H4A3J1, Canada; Metabolic Disorders and Complications program, Research Institute of the McGill University Health Centre, Montréal, QC H4A3J1, Canada; Department of Pharmacology and Therapeutics, McGill University, Montréal, QC H3G1Y6, Canada

**Author notes:** Corresponding author: Dr. Shafaat A. Rabbani, Research Institute of the McGill University Health Centre, 1001 Décarie Blvd. (Glen site), Montréal, QC H4A3J1, CANADA.

**Keywords:** DNA methylation readers, epigenome, breast cancer, metastasis, MBD

## Abstract

Methyl-CpG-binding domain protein 2 (Mbd2), a reader of DNA-methylation, has been implicated in the progression of several types of malignancies, including breast cancer. To test whether Mbd2, which is overexpressed in human breast cancer samples and in MMTV-PyMT mammary pads, plays a causal role in mammary tumor growth and metastasis we depleted Mbd2 in transgenic MMTV-PyMT model of breast cancer by cross-breeding with Mbd2 knockout mice to generate heterozygous (PyMT;*Mbd2^+/-^*) and homozygous (PyMT;*Mbd2^-/-^*) animals. We found that *Mbd2* depletion caused a gene dose-dependent delay in mammary tumor formation, reduced primary tumor burden, and lung metastasis at the experimental endpoint. In addition, animals from the PyMT;*Mbd2^-/-^* group survived significantly longer compared to the wildtype (PyMT;*Mbd2^+/+^*) and PyMT;*Mbd2^+/-^* arms. Transcriptomic and proteomic analyses of the primary tumors obtained from PyMT;*Mbd2^+/+^* and PyMT;*Mbd2^+/-^* groups revealed that *Mbd2* depletion alters several key determinants of the molecular signaling networks related to tumorigenesis and metastasis, which thereby demonstrate that Mbd2 is regulating transcriptional programs critical for breast cancer. To our knowledge, this is the first study demonstrating a causal role for a DNA-methylation reader in breast cancer. Results from this study will provide the rationale for further development of first-in-class targeted epigenetic therapies against Mbd2 to inhibit the progression of breast and other common cancers.

## Introduction

DNA methylation is an evolutionarily ancient epigenetic process which, through the modulation of chromatin structure, regulates gene expression in a context-dependent manner (1, 2). The chemically reversible process of DNA methylation is mediated by a family of “writer” enzymes known as DNA methyltransferases (DNMTs) that catalyze the addition of methyl-moieties to the appropriate bases on the genome (3). A family of evolutionarily conserved “reader” proteins known as methyl-binding proteins (MBPs) then recognize, interpret, and relay the information from these methylation marks into different gene regulatory functionalities (2).

Aberrant DNA methylation is recognized as a paradigmatic hallmark of human cancer (4), which led to its identification as an attractive therapeutic target. The first-generation epigenetic drugs developed to target the methylome are DNMT inhibitors (Vidaza and Decitabine) that showed robust clinical utilities against several hematological malignancies (5). At the molecular level, through non-specific global demethylation, the DNMT inhibitors not only relieves the transcriptional repression of critical tumor-suppressors (6) but also cause transcriptional activation of several known prometastatic genes (7). Moreover, these drugs are highly toxic, less bioavailable, and showed a modest anti-cancer response in case of solid tumors (8), which opens up novel avenues to selectively target the MBPs as an alternative approach to reverse the DNA-methylation mediated epigenetic abnormalities in cancer.

Among the different MBPs, methyl-CpG-binding domain protein 2 (MBD2) is positioned as a suitable anti-cancer drug target since its expression is deregulated in several human malignancies (2) and shows a relatively higher affinity to bind to the methylated DNA near the promoters of various known tumor suppressor genes to cause their transcriptional repression (9). Moreover, genetic knockout (KO) of the *Mbd2* gene in mice produced viable off-springs, which suggests that the gene is not required for maintaining standard physiological functions (10) and, thus, could be used for targeted epigenetic therapies. Indeed, genetic depletion of *Mbd2* has been shown to protect mice from developing intestinal (11) and lymphoid malignancies (12). MBD2 also plays an important role in immune regulation by upregulating the expression of forkhead box P3 (*Foxp3*), which is the marker for regulatory T cells (Treg) (13). Other studies have shown that the immunosuppressive Treg cells infiltrate into the tumors to promote immune evasion and pro-tumorigenic microenvironment and are associated with poor cancer prognosis in patients (14, 15).

The role of MBD2 in breast cancer progression thus far has been studied through gene knockdown *in vitro* and subsequent implantation into xenograft models where *Mbd2* depletion has shown potent anti-cancer effects through inhibition and hypermethylation of prometastatic genes (16, 17). However, these models lack functional immune systems and are not capable of fully reflecting the exact role of MBD2 during the highly complex multistep progression of human breast tumors. To dissect the role of MBD2 in a mouse model with a relatively faithful representation of breast tumor progression, we used a transgenic MMTV-PyMT (mouse mammary tumor virus-polyoma middle tumor-antigen) model where the spontaneous and pregnancy-independent expression of PyMT oncoprotein results in the synchronous appearance of multifocal breast tumors that metastasize predominantly to the lung (18, 19). Although the PyMT oncoprotein is not present in human breast tumors and other triggers initiate breast cancer, the step-wise progression of the murine mammary tumor from benign premalignant stage to a highly malignant invasive stage as well as the activation of downstream molecular signaling pathways resemble that of human breast cancer progression (20). This positions the PyMT model as a system to assess the oncogenic function of a particular gene in the evolution of breast malignant transformation in a whole organism (21).

Herein, using a gene-knockout based molecular genetics approach, we demonstrate that a reduction in the ability to read and interpret epigenetic modification due to the depletion of *Mbd2* gene significantly decrease mammary tumor burden and metastasis in transgenic MMTV-PyMT model of breast cancer.

## Results

### *Mbd2* KO attenuates primary breast tumor growth and metastases in MMTV-PyMT

We first interrogated the publicly available proteomics datasets from the Clinical Proteomic Tumor Analysis Consortium (CPTAC) to check the expression of MBD2 and found MBD2 expression is significantly upregulated in different subtypes of human breast tumors compared to their normal counterpart (Figure 1A). Interestingly, Mbd2 expression is upregulated in PyMT mammary fat pads as compared with the fat pad of wildtype C57BL/6 and PyMT animals suggesting that Mbd2 upregulation by the *PyMT* gene precedes malignant transformation consistent with the hypothesis that Mbd2 is mediating some of the oncogenic effects of pyMT (Figure 1B). Mbd2 expression increased as the tumor progressed to an advanced stage at week 20 compared to the fat pads collected at a pre-cancerous stage on week 11 (Figure 1B).

**Figure 1:**
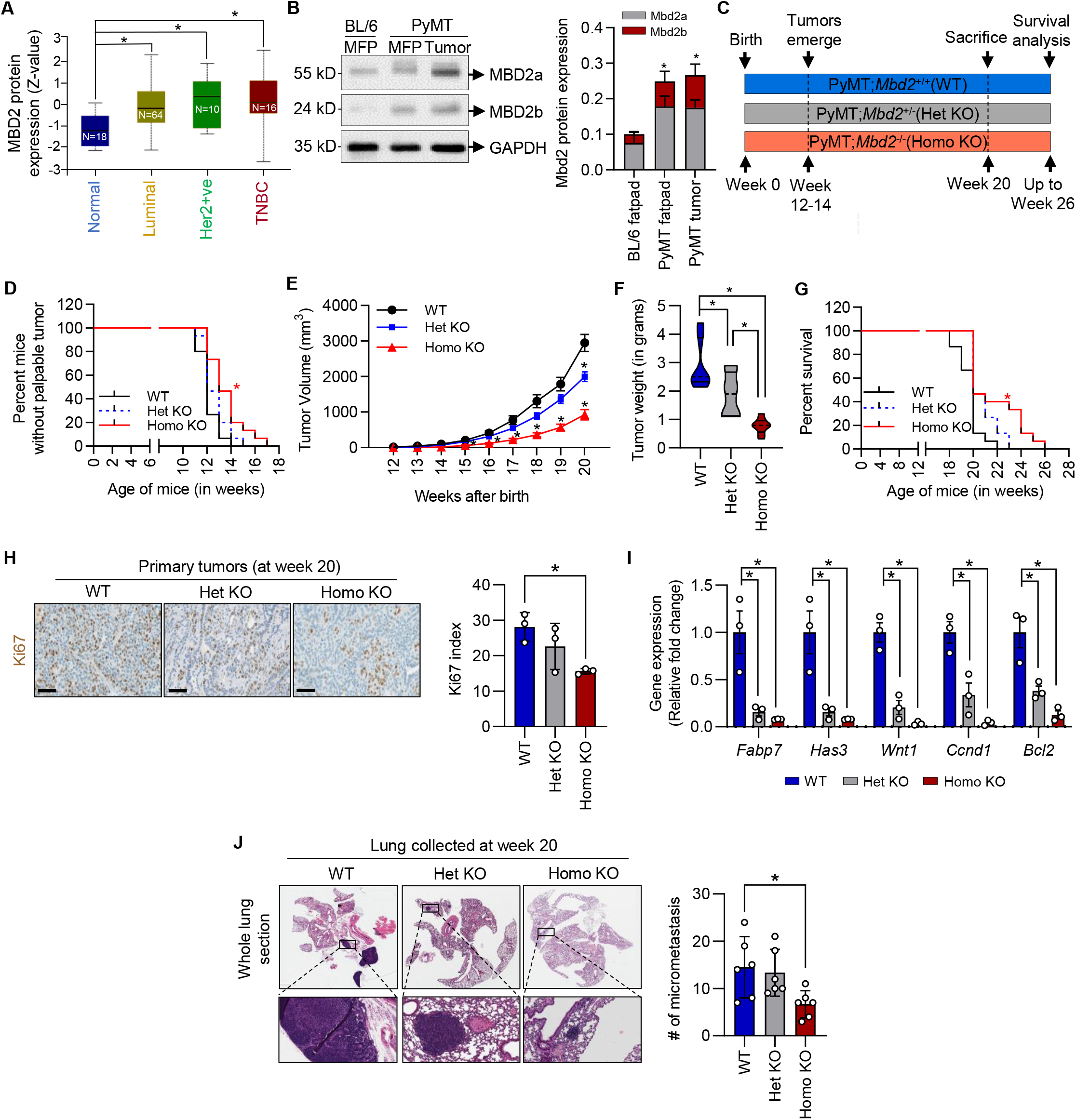
MBD2 is upregulated in breast tumors, and knockout of the *Mbd2* gene affects tumor and progression in transgenic PyMT mice. **(A)** MBD2 protein levels in normal subjects and different subtypes of breast cancer. **(B)** Immunoblot of the mouse Mbd2 protein from lysates obtained from mammary fat pad of 11-week-old female C57BL/6 (lane 1) and PyMT mice (lane 2), and mammary tumors 20-week-old PyMT mice (lane 3). GAPDH was used as a loading control. The right panel shows normalized densitometric quantification of the total Mbd2 signals (n=3 animals/group). **(C)** Schematic representation of the endpoints of this study. **(D)** A Kaplan-Meier curve showing tumor emergence in the three groups. **(E)** Tumor volumes were measured at weekly intervals until sacrifice (n=15 animals/group). **(F)** Violin plot showing the distribution of tumor weight in each group measured at week 20. At least eight animals in each group were sacrificed at this point in addition to the ones that researched to a level requiring humane sacrifice. The rest of the animals were used for Kaplan-Meier survival analysis in **(G)** until they required humane sacrifice. **(H)** Immunohistochemical staining of the primary tumors from each group of animals with an antibody against Ki67 proliferation marker (left panel). The percentage of Ki67 positive tumor cells was determined from five high-power fields for each sample and plotted as a bar graph in the right panel (n=3 animals/group). **(I)** qPCR of several known cancer-related genes using RNA extracted from primary tumors (n=3 animals/group). **(J)** H&E staining of formalin-fixed lung tissues obtained at experimental endpoint at week 20. The number of micrometastases were counted and plotted as a bar graph (n=6 animals/group). Results are represented as the mean ± SEM, and statistically significant differences were determined using ANOVA followed by *post hoc* Tukey’s test. *P* < 0.05.

Next, to test whether *Mbd2* plays a causal role in tumorigenesis, we generated female PyMT;*Mbd2^+/-^* (heterozygous KO) and PyMT;*Mbd2^-/-^* (homozygous KO) mice in C57BL/6 background using a cross-breeding strategy, and compared their tumor growth kinetics with PyMT;*Mbd2^+/+^* (denoted as wildtype hereafter) animals from week 11 after birth until sacrifice (Figure 1C). As shown by the Kaplan-Meier curve in Figure 1D, animals from the wildtype group started to develop palpable tumors at around week 11 (day 77) after birth, and by week 14 (day 98) of age, all animals from the wildtype group had developed primary tumors [wildtype tumor incidence: 77–98 days; 50% mice with palpable tumor, T_50_=84 days] (Figure 1D). The onset of palpable tumor was not significantly different in case of the animals from the heterozygous KO group where primary mammary tumors started to emerge at around the same time as control i.e. by week 11, and by week 15 (day 105) of age, 100% of the animals in this group showed palpable mammary tumors [heterozygous KO group tumor incidence: 77–105 days; T_50_=84 days]. In the homozygous KO group, tumors started to emerge from week 12 (day 84), and all the animals developed palpable tumors by week 16 (day 112) of age [homozygous KO tumor incidence: 84– 112 days; T_50_=91 days]. The onset of the palpable tumor was significantly delayed in homozygous KO group compared to the wildtype arm (log rank *P*=0.002), suggesting the possible involvement of *Mbd2* in mammary tumor onset in this model.

Next, we compared the tumor growth kinetics of the animals from the three groups by measuring the primary tumor volumes at weekly intervals from the time the animals had started to develop measurable tumors until they were sacrificed. The experimental endpoint was set at week 20 after birth when most of the wildtype mice reached the humane endpoint. We found that the tumor growth over time was significantly reduced in a gene dose-dependent manner (Figure 1E),, an observation consistent with similar results shown in case of intestinal tumorigenesis (11). We then measured the total weight of the extirpated tumors of the animals scarified either before or at week 20 after birth and found a significant reduction in tumor weight in the heterozygous and homozygous KO groups in comparison with the wildtype animals (Figure 1F). To test whether *Mbd2* plays a role in prolonging the survival of the mammary tumor-affected mice, we kept several animals from each group beyond the experimental end point at week 20 and found that the animals from the homozygous KO group reached the tumor volume requiring humane sacrifice significantly later than animals from wildtype and heterozygous KO groups (Figure 1G).

We then evaluated the formalin-fixed mammary tumor tissues from each group with an antibody for the Ki67 cell proliferation marker and found a significant decrease of Ki67 positive cells in homozygous KO group (Figure 1H). Next, using the RNA from flash frozen lung tissues collected at around week 20, we checked for the expression of several known cancer genes (*Fabp7, Has3, Wnt1, Ccdn1, Bcl2*) that were previously shown to be regulated either directly through MBD2 or its downstream signaling pathways, and found a significant decrease in their expression in the homozygous KO tumors compared to the tumors from the wildtype animals (Figure 1I).

The female MMTV-PyMT animals develop pulmonary metastases which allowed us to examine the role of Mbd2 in metastatic dissemination of primary breast tumor cells into distant secondary organs, a condition that is common in clinical settings in breast cancer patients. For that, we used the formalin fixed lung tissue sections collected at sacrifice at week 20, stained them with H&E and found a significant decrease in the number of micrometastases in the animals from homozygous KO group in comparison with the wildtype and heterozygous KO groups (Figure 1J).

Taken together, these results demonstrate that the genetic ablation of the *Mbd2* gene significantly attenuates the ability of the MMTV-PyMT animals to grow spontaneous mammary tumors with significant reduction in pulmonary metastasis.

### Mbd2 depletion affects the several crucial players of PyMT-mediated oncogenic signaling pathways

PyMT is a membrane-associated oncoprotein that does not have any kinase activity of its own (21). However, when it interacts with receptor tyrosine kinases (RTKs) [for example, c-Src, p85 subunit of phosphoinositide 3-kinase (PI3K)], the resultant protein complex gains constitutive tyrosine kinase activities required for activation of downstream signaling pathways to promote cellular transformation, growth, and survival (18). To examine whether the Mbd2 depletion directly impairs the crucial signaling molecules in PyMT mediated tumorigenesis, we first checked the levels of activated c-Src, PI3K, and AKT in proteins extracted from wildtype and *Mbd2* KO tumors.

Towards these goals, we first compared the level of c-Src phosphorylation (p-c-Src) at Y416 residue, which is required for the oncogenic transformation function of the PyMT-c-Src complex (22), and found a substantial decrease in the total c-Src levels in *Mbd2* KO tumors which cause a net decrease in p-c-Src (Y416) level (Figure 2A). Previous studies have shown that PyMT also complexes with the p85 subunit of PI3K to mediate transformation (23, 24). When we checked the phosphorylation status of PI3K (at Y458), a significant decrease was observed in *Mbd2* KO tumors (Figure 2A). The total PI3K level did not change between wildtype and *Mbd2* KO tumors (Figure 2B). Moreover, a significant impairment in the activation of AKT (at S473), which is a downstream signaling molecule of the PI3K signaling pathway, was observed in *Mbd2* KO tumors compared to their wildtype counterparts (Figure 2A-B).

**Figure 2:**
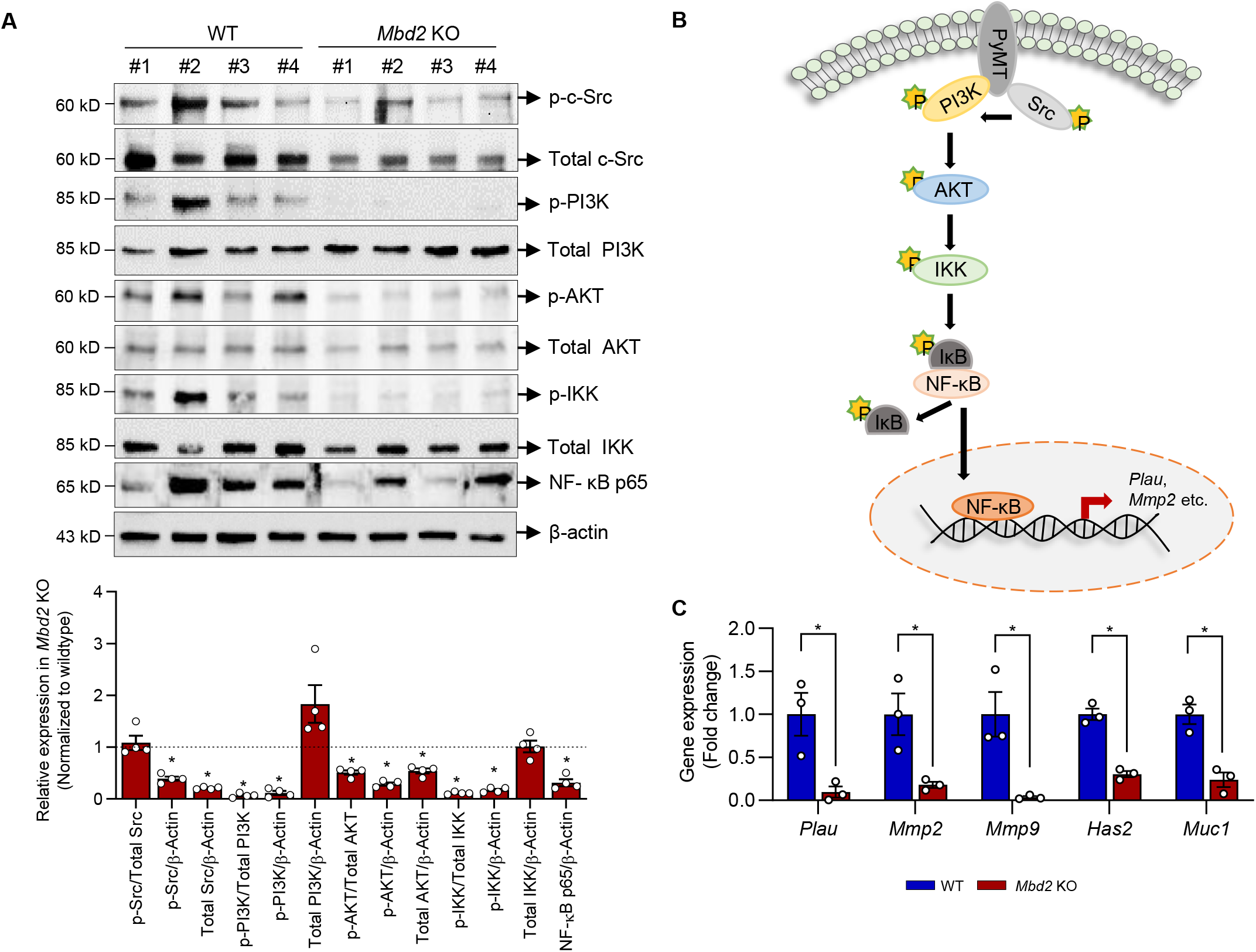
*Mbd2* depletion interferes with the PyMT-dependent activation of the PI3K/Akt/NF-κB axis. **(A)** Immunoblots from whole tumor lysates obtained from four distinct animals/group (upper panel). Densitometric quantification of the bands for each protein was determined and plotted as mean ± SEM (n=4 tumors/group) (lower panel). **(B)** Schematic diagram of PyMT-mediated oncogenic signaling pathways downregulated by Mbd2 depletion. **(C)** qPCR of several NF-κB regulated genes (n=3/group). Statistical significance was determined using the student’s *t*-test. \**P* < 0.05.

Since AKT is known to activate IKK activity through phosphorylation and thereby activate the downstream NF-κB signaling cascade (25), we checked and found that the phosphorylation of IKK and the levels of activated NF-κB (p65 subunit) are decreased in the *Mbd2* KO tumors compared wildtype (Figure 2A-B). Furthermore, several known target genes of NF-κB transcription factor like *Plau* (also known as *uPA*), *Mmp2*, *Mmp9*, *Has2,* and *Muc1* were downregulated in the RNA extracted from *Mbd2* KO tumors (Figure 2C). Taken together, these data suggest that *Mbd2* depletion impairs the ability of the PyMT oncoprotein to stimulate the oncogenic PI3K/Akt/NF-κB axis.

### Transcriptome and proteome analyses reveal decrease in EMT markers in *Mbd2* KO tumors

We then investigated the transcriptomic changes triggered by the genetic depletion of *Mbd2* in primary tumors by RNA-seq (n=3 samples/group). We found that, in comparison to the wildtype arm, the primary tumors from the Mbd2 KO group caused a significant change in the expression of 453 genes (|log2(FoldChange)| > 1, *P*<0.05). The expression changes occurred in both directions i.e., 121 genes were upregulated, and 332 genes were downregulated in the *Mbd2* KO tumors (Figure 3A). Using the Ingenuity Pathway Analysis (IPA) tool (26), we found that some of notable ‘molecular and cellular functions’ enriched with differentially expressed genes (DEGs) in *Mbd2* KO tumors include cellular movement, cell-cell signaling & interaction, cellular assembly & organization, molecular transport, cell death & survival, and cellular growth & proliferation (Supplementary Figure S1A). IPA analyses also identified that the top two ‘disease and disorders’ pathways significantly enriched with DEGs in *Mbd2* KO tumors include ‘cancer’ and ‘organismal injury and abnormalities’ (Supplementary Figure S1B). We then subjected the entire list of significant DEGs to pathway analyses using the Pathway Interaction Database (PID) and found a statistically significant enrichment of the genes in integrin signaling, syndecan-4 mediated signaling, adherens junction stability and disassembly pathways (Figure 3B). Interestingly, most of these pathways are related to cancer progression, especially epithelial-to-mesenchymal transition (EMT). Therefore, we next overlapped the EMT related genes from the publicly available dbEMT database with genes downregulated in *Mbd2* KO tumors and found a significant intersection of 35 genes (hypergeometric test, p<0.05, Figure 3C). A heatmap of the differentially expressed EMT genes in *Mbd2* KO tumors is shown in Figure 3D. To validate the results from the RNA-Seq experiment, we performed a quantitative polymerase chain reaction (qPCR) analysis of several crucial EMT genes (*Sparc, Spp1,* and *Cdh2*) and found a concordant decrease in their expression in the *Mbd2* KO tumors (Figure 3E). Importantly, *Sparc, Spp1,* and *Cdh2* are all upregulated in the human breast cancer tumors from the TCGA database (Figure 3F), suggesting the clinical relevance of MBD2 regulated genes in breast cancer. We also performed qPCR validation of several known tumor suppressor genes (*Brca1, Dusp5*) that were upregulated upon *Mbd2* gene KO (Supplementary Figure S1C).

**Figure 3:**
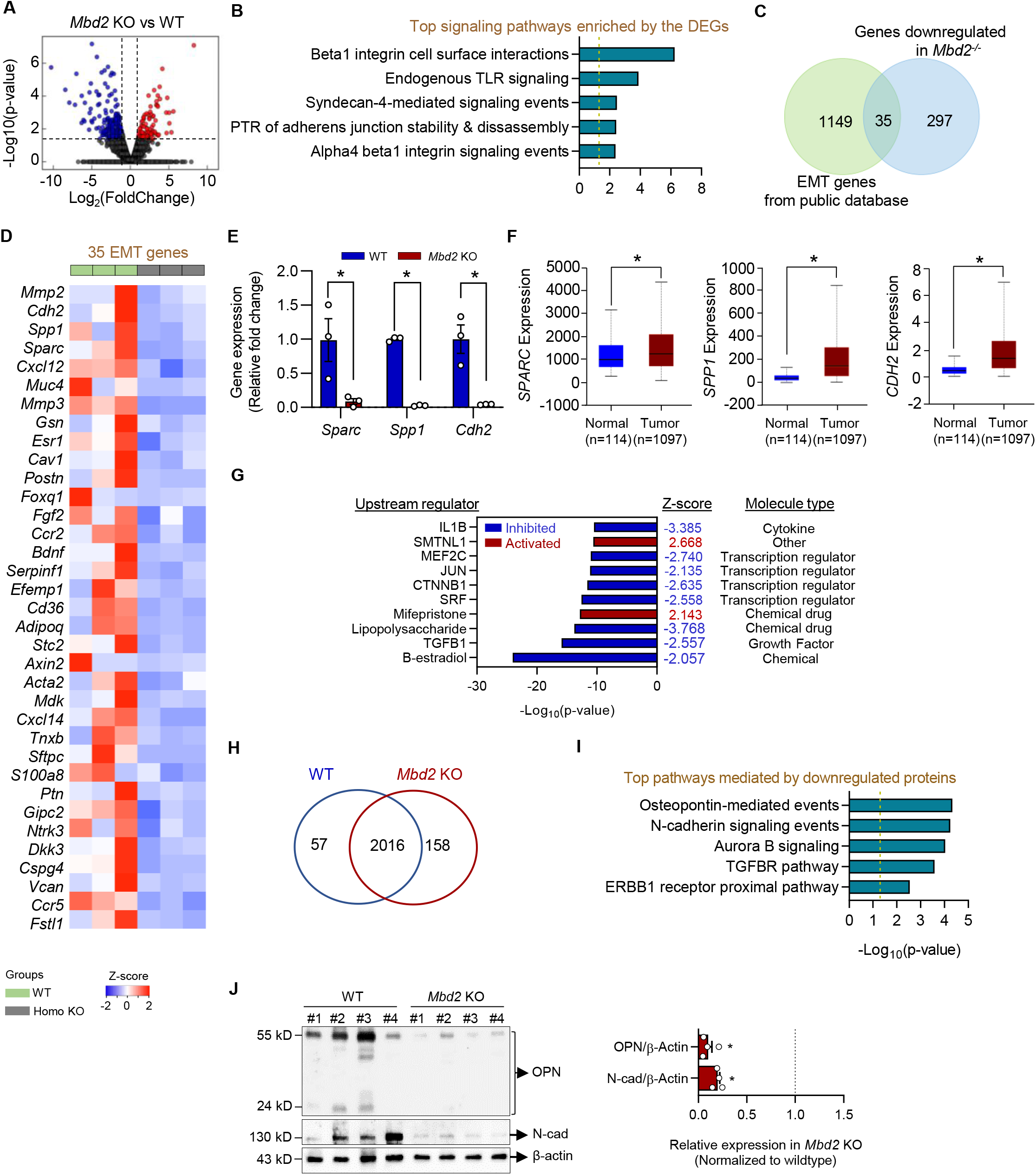
Transcriptomic and proteomic analyses of wildtype and *Mbd2* KO tumors. **(A)** RNA extracted from mammary tumors of control and knockout mouse were subjected to RNA-Sequencing (n=3 samples/group). A volcano plot showing the DEGs (332 downregulated and 121 upregulated) in the Mbd2 KO vs. wildtype groups. **(B)** Pathway enrichment analysis (PID database) of the DEGs in *Mbd2* KO tumors. **(C)** Venn diagram of the 332 downregulated genes in *Mbd2* KO tumors with the complete repertoire of genes from the epithelial–mesenchymal transition (EMT) database showed an overlap of 35 genes that are presented in the heatmap in **(D)**. **(E)** qPCR validation of the selected EMT-related genes (*Sparc, Spp1, Cdh2*) obtained from RNA-Seq was done using tumoral RNA from at least three animals/group. **(F)** The gene expression pattern of the human orthologs of the qPCR validated genes in normal and breast tumors according to the TCGA database. **(G)** Top ten upstream regulators of the DEGs obtained from RNA-Seq as predicted by IPA upstream regulator analyses tool. **(H)** Venn diagram of the common and unique protein hits in wildtype and *Mbd2* KO animals according to the proteomics analyses. **(I)** Pathway enrichment analysis (PID database) of the significantly downregulated proteins in *Mbd2* KO tumors. **(J)** Analysis of Osteopontin and N-Cadherin proteins in the immunoblots of whole tumor lysates. The right panel shows the densitometric quantification of the immunoblots plotted as mean ± SEM (n=4 tumors/group). Statistical significance was determined using the student’s *t*-test. \**P* < 0.05.

Next, we employed the IPA upstream regulator analysis (URA) for the identification of potential transcription factors, cytokines, growth factors, or any chemical entities that could regulate the gene expression changes seen in our transcriptome datasets (Figure 3G). Interestingly, the top upstream regulators that are predicted to be downregulated include TGFB1, SRF, CTNNB1, and JUN, all of which are known to be involved in the EMT pathway. In addition, we employed an alternative approach for predicting the upstream transcription factors of the DEGs identified in *Mbd2* KO tumors by searching the ChIP Enrichment Analysis (ChEA) database (27). This analysis identified 57 transcription factors that include several EMT-related transcription factors like MTF2, SOX2, SOX9, JUN and others (Supplementary Figure S1D).

We next performed a uHPLC/MS-MS to assess the proteomic differences between the mammary tumors extracted from wildtype and *Mbd2* homozygous KO animals. A total of 2231 proteins were identified. To gain insights into the biological processes affected by the deletion of *Mbd2* gene, we focused on the changes in the abundance of proteins between the two groups as a measure of differential protein expression. This approach identified 215 proteins with differential abundance, of which 158 are upregulated and 57 are downregulated in the KO samples compared to the wildtype counterparts (Figure 3H). Protein-protein interaction meta-analyses of the 215 proteins revealed their involvement in biological processes like metabolism of RNA, telomerase RNA localization, cytoplasmic translation initiation. Since our transcriptome analyses revealed the downregulation of several key cancer-related genes in homozygous KO tumors, we then focused on the downregulated proteins. Pathway analysis of the downregulated proteins in homozygous KO tumors revealed their involvement in osteopontin-mediated signaling, N-cadherin signaling, TGFBR pathways, all of which are involved in the EMT (Figure 3I). We next validated the downregulation of Osteopontin (encoded by *Spp1* gene) and N-cadherin (encoded by *Cdh2* gene) in protein level (Figure 3J), which were also downregulated at the RNA level. Taken together, these observations indicate the possible involvement of Mbd2 in modulating different components of EMT during breast tumor progression in this model.

We also overlapped the list of differentially expressed proteins and RNAs obtained from our proteomics and transcriptomics studies and found an overlap of 10 entities, all of which showed concordant expression patterns (Supplementary Figure 1E-F).

### Mbd2 modulates the oncogenic Pvt1-Myc axis

Emerging evidence suggests that pervasive transcription of genes beyond the proteincoding boundaries of the genome plays a diverse array of functions in the regulation of gene expression as well as the pathogenesis of disease (28–30). Therefore, we next focused on deciphering the lncRNAs that are differentially expressed upon homozygous deletion of *Mbd2* gene using our RNA-Seq data and found that 60 lncRNAs are differentially expressed in PyMT;*Mbd2^-/-^* primary tumors compared to that of PyMT;*Mbd2^+/+^* (Supplementary Figure S2A). Gene ontology (GO) analyses of the lncRNAs revealed their involvement in a wide range of biological process related to RNA processing and transcriptional regulation (Supplementary Figure S2B). Since a vast majority of the lncRNAs are still uncharacterized in the context of cancer, we focused on the lncRNAs with known biological function(s) or known human orthologs in the TCGA breast cancer patient dataset. A curated list of 16 lncRNAs that fulfill these criteria is shown as heatmap in Figure 4A. We further validated the expression of several lncRNAs [X-inactive specific transcript (*Xist*), chaperonin containing Tcp1 and subunit 6a (*Cct6a*), and plasmacytoma variant translocation 1 (*Pvt1*)] by qPCR (Figure 4B), where the expression change showed concordance with the RNA-Seq results. The expression of the corresponding orthologous genes in human breast cancer patients from TCGA database showed that *XIST* expression is significantly downregulated while *CCT6A,* and *PVT1* expressions are significantly upregulated (Figure 4C). The *PVT1* and *MYC* genes are coexpressed in human genome when we performed a Pearson correlation analysis using data obtained from 4307 patients (Figure 4D). Similar to their human counterpart, these two genes are also located in close proximity on the chromosome 15 of mouse genome (Figure 4E). We, therefore, checked whether *Myc* is differentially regulated in the PyMT;*Mbd2^-/-^* tumors and found a significant repression of its expression both in the transcript and protein levels in tumors obtained from PyMT;*Mbd2^-/-^* compared to that of PyMT;*Mbd2^+/+^* (Figure 4F-G). These results suggest the possible role of Mbd2 in modulating the oncogenic Pvt1-Myc axis in cancer.

**Figure 4:**
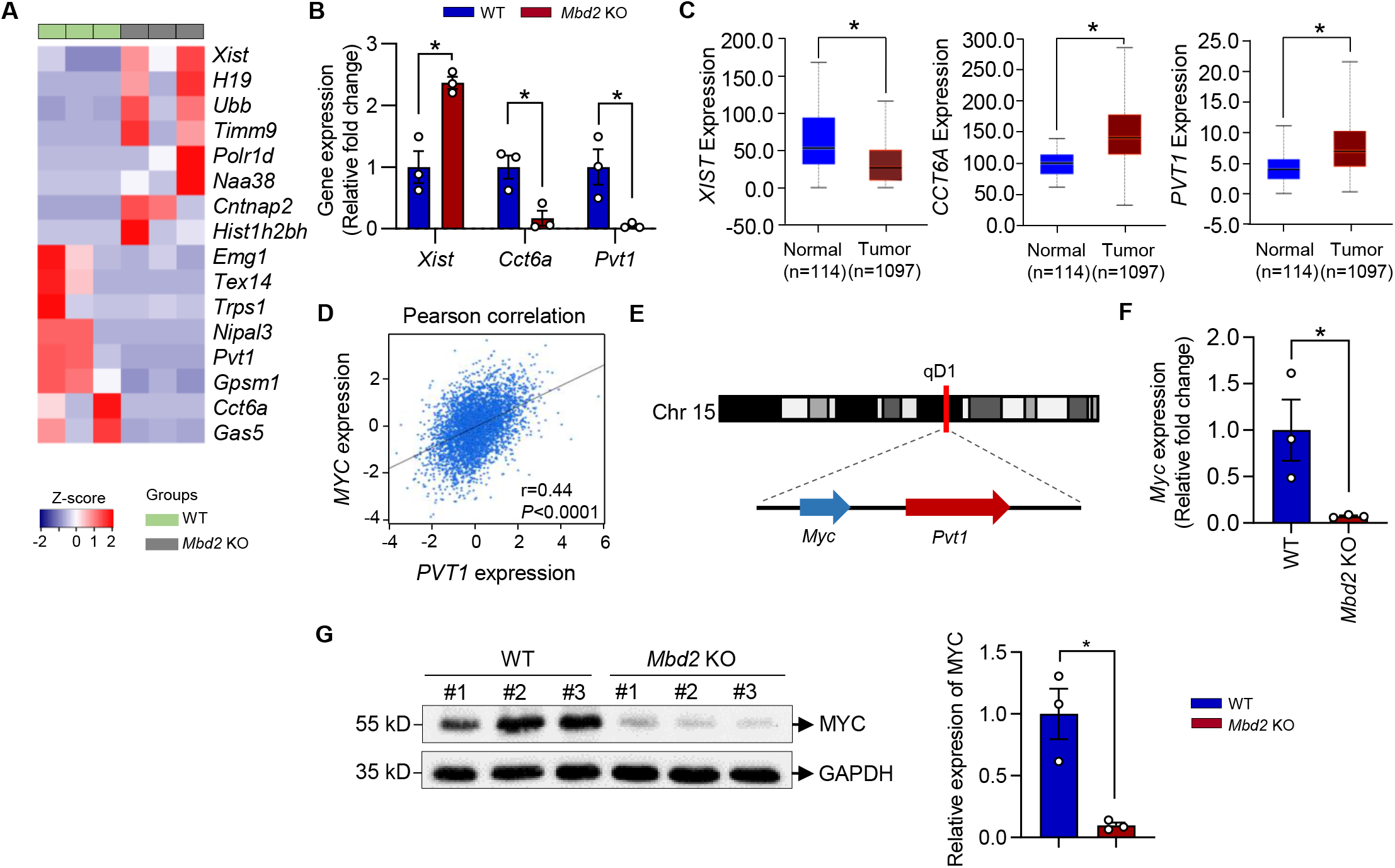
Analyses of differentially expressed lncRNAs in *Mbd2* KO tumors. **(A)** Heatmap of differentially expressed lncRNAs in *Mbd2* KO tumors with known biological function(s) or known human orthologs in the TCGA breast cancer patient dataset. **(B)** qPCR validation of the selected lncRNAs (*Xist, Cct6a, Pvt1*) obtained from RNA-Seq was done using tumoral RNA from at least three animals/group. **(C)** The gene expression pattern of the human orthologs of the qPCR validated genes in normal and breast tumors according to the TCGA database. **(D)** Pearson correlation between human *PVT1* and *MYC* genes using data obtained from 4307 patients in TCGA, GSE81538, and GSE96058. **(E)** Schematic of the chromosomal location of *Myc* and *Pvt1* genes revealing proximity of the genes on the mouse genome (not drawn to scale). **(F-G)** RNA and protein levels of Myc is significantly altered in *Mbd2* KO tumors relative to wildtype counterparts. Results are shown as mean ± SEM (n=3 tumors/group). Statistical significance was determined using the student’s *t*-test. \**P* < 0.05.

## Discussion

Mbd2 has been previously suggested to be involved in promoting breast and other cancers growth and metastasis, however past studies were performed in cancer cell lines and xenografts in mice and used either antisense or siRNA knockdown approaches. Since these methodologies are highly confounded, the main question remained as to whether Mbd2 plays a causal role in breast cancer. In addition to the general well-known limitations of cell culture experiments and the epigenomic changes that occur in cultured cells, these studies could only address the role of Mbd2 once cancer has evolved but did not capture its role during premalignant and early stages. Moreover, the studies do not capture the context of normal organismal development from birth to adulthood and the complex tumor-host relationships that occur in a whole animal including the emerging significance of the immune system. Xenograft studies involve the use of severely immunodeficient mouse strains to avoid rejection of the graft by the host and therefore pass over the role of immune system in cancer progression.

In this study, we genetically depleted *Mbd2* in the well-characterized transgenic MMTV-PyMT mouse model that recapitulates the stepwise progression of tumors from localized to metastatic variant and share similar pathology and biomarkers found in patients with metastatic breast cancer (20). Since the gene is depleted preconception, gene loss certainly precedes tumor initiation and thus this study design provides definitive conclusions on the causal role of *Mbd2* in breast cancer. Our data demonstrate that the PyMT oncogene driven spontaneous mammary tumor formation is delayed upon homozygous depletion of the *Mbd2* gene. *Mbd2* might not be involved in tumor initiation as tumors are delayed but nevertheless appear in the KO but it is required for tumor growth since tumor growth is significantly delayed by *Mbd2* deficiency. The late appearance of tumors in *Mbd2*-/- mice might be a consequence of slowing down tumor growth and therefore tumors reach detection size later. This conclusion is further supported by the significant decrease of Ki67 positive cells in homozygous KO group suggesting reduced proliferation of the KO cancer cells. This observation is different than a previous study where *Mbd2* depletion protected mice from the development of intestinal cancer and reduced the number of polyps suggesting a role in tumor initiation in intestinal cancer (11). Moreover, we found that even at a haploinsufficiency state where one of allele is not present, the heterozygous PyMT;*Mbd2^+/-^* group is still able to cause a significant reduction in primary breast tumor burden further implicating *Mbd2* in tumor growth.In addition to tumor growth, lung metastasis is significantly reduced upon the genetic ablation of the *Mbd2* gene (Figure 1). This is consistent with previous results implicating *Mbd2* in breast cancer metastasis (31, 32); our results provide evidence that *Mbd2* plays a causal role in breast cancer metastasis. These results indicate a multifaceted role of Mbd2 during mammary tumor progression.

Mbd2 is upregulated by PyMT and upregulation of Mbd2 precedes tumor formation consistent with the idea that Mbd2 is mediating the effects of PyMT on cellular pathways that precipitate uncontrolled growth and metastasis; Mbd2 depletion blocks activation of these pathways. PyMT activate a panel of downstream oncogenic signaling pathways (for example, Src, PI3K/Akt) that are involved in tumor cell proliferation, survival, inhibition of apoptosis and metastasis (21). Our analyses of the proteins obtained from the primary tumors revealed that *Mbd2* deletion represses the activation of PI3K/Akt axis (Figure 2), which has been previously shown to be essential for mammary tumorigenesis (33). Furthermore, *Mbd2* depletion interferes with the oncogenic NF-κB signaling pathway which results in the repression of critical cancer-related downstream targets of NF-κB like *Plau, Mmp2, Mmp9* and several others. These targets are involved as well in breast cancer invasion and metastasis. It is unclear how Mbd2 alters signaling cascades since it seems to affect phosphorylation but not the level of the proteins. This is intriguing since Mbd2 is a well-established regulator of gene expression. Is it possible that Mbd2 has an additional role in signaling? Alternatively, it might regulate signaling cascade through regulation of genes that control the signaling events described in our study. This question needs to be addressed in future studies.

Transcriptome analyses provide insights into the global gene expression changes mediated by *Mbd2* KO. Although Mbd2 is generally believed to be a suppressor of gene activity through recruitment of the Nurd complex and histone deacetylase to methylated promoters (34). and a previous study in an isogenic model of breast cancer transformation showed that the majority of genes whose expression changes following *Mbd2* knockdown are downregulated(35), we show here that the majority of genes are silenced rather than activated by *Mbd2* depletion. Many of these genes play an active role in cancer growth and metastasis. Thus, Mbd2 serves as an activator of several cancer genes in our model. However, although the silencing activity of Mbd2 has been emphasized in past studies, several studies have shown that Mbd2 is involved in gene activation as well. For example, Baubec et al., show that Mbd2 plays a bimodal role in embryonal stem cells and that Mbd2 binds to methylated regions in the genome which are enriched with repressive marks as well as to active unmethylated regulatory regions that are enriched with DNAse hypersensitive sites and activating chromatin marks (36). Several studies have shown that Mbd2 activates promoters and induces hypomethylation (37–44) (45). Relevant to our studies, we have previously shown that depletion of Mbd2 leads to silencing and hypermethylation of prometastatic genes in breast cancer cells(31, 32). Mbd2 is most probably targeted by different factors to either activate or silence gene expression. It was proposed that the NurD complex targets Mbd2 to active unmethylated regions in embryonal stem cells (36). Stefanska et al., has shown that Mbd2 activates and triggers hypomethylation of cancer promoting genes in liver cancer(46) and that the transcription factor CCAAT/enhancer-binding protein alpha (CEBPA)CEBPA recruits Mbd2 to its targets to trigger transcription onset (43). Mbd2 is recruited to the *Foxp3* gene in regulatory T cells which results in loss of methylation possibly through recruitment of Tet2(45).Thus, our data is consistent with previous data suggesting that Mbd2 plays a role in activation of several cancer related gene pathways, most probably through recruitment to specific genomic regions by transcription factors that are activated by triggers of cancer such as PyMT. Further studies are required to test this hypothesis. Our transcriptome analyses supports a bimodal role for Mbd2 in breast cancer both activating genes that promote and silencing genes that suppress cancer.Importantly, our RNA-Seq results indicated that several key EMT pathway related gene signatures encoding for mesenchymal markers like N-Cadherin, Osteopontin were significantly downregulated by Mbd2. EMT is a highly dynamic biologic process that directs the epithelial cells within a particular tissue to go through multiple biochemical changes rendering their transformation to more invasive mesenchymal cells (47). Therefore, a decrease in the expression of the mesenchymal marker reduces the metastatic spread of primary tumors. A proteomic analysis of the lysates obtained from wildtype and *Mbd2* KO tumors further indicated and confirmed that the top pathways enriched by the differentially downregulated proteins in *Mbd2* KO tumors are Osteopontin, N-cadherin and TGFR mediated signaling events-all of which involved in EMT (Figure 3J).

To our knowledge, this is the first report describing multifaceted function of an epigenetic reader during mammary tumor progression and metastasis. Since *Mbd2* depleted animals are viable and fecund, it would be similarly interesting to test whether targeting Mbd2 and/or its methylated DNA binding ability would produce a similar anti-cancer effect in patients with breast cancer to reduce cancer-associated morbidity and mortality. It will also open a novel avenue for the use of targeted epigenetic therapies against methylation abnormalities in cancer as single agent monotherapy or in combination with standard of care treatment regimens.

## Materials and Methods

### Mouse strains and genotyping

The embryo straws for the *Mbd2* heterozygous KO mice (*Mbd2*^+/-^) in C57BL/6 background were a kind gift by Dr. Brian Hendrich (Department of Biochemistry, University of Cambridge). The embryos were first recovered from the straws at the Transgenic Core Facility, Rosalind and Morris Goodman Cancer Research Centre, McGill University and then they were implanted into two female foster mice to generate the heterozygous KOs (*Mbd2*^+/-^). Afterwards, the animals were bred and maintained at the Animal Resource Division of the Research Institute of the McGill University Health Center. The breeder pair of male heterozygous MMTV-PyMT (in C57BL/6 background) carrying a single copy of the PyMT transgene and noncarrier wildtype C57BL/6 female were purchased from The Jackson Laboratory (Bar Harbor, ME, USA) and they were crossed to generate MMTV-PyMT (also denoted as PyMT) mice that are wildtype for the *Mbd2* gene. The male PyMT mice were then crossed with female *Mbd2^+^*^/-^ mice to generate the PyMT;*Mbd2*^+/-^ (heterozygous KO for *Mbd2*) littermates. Next, male PyMT;*Mbd2^+/-^* and female *Mbd2^+/-^* mice were crossed to generate female PyMT;*Mbd2^+/+^*, PyMT;*Mbd2^+/-^*, and PyMT;*Mbd2^-/-^* (homozygous KO for *Mbd2*) for assessing tumors. All animals were heterozygous for the PyMT transgene.

For genotyping, DNA was extracted from the tails snips obtained from the mice followed by polymerase chain reaction (PCR) using primers for the detection of specific alleles. The PyMT allele was detected using the following primers: GGAAGCAAGTACTTCACAAGGG (forward) and GGAAGTCACTAGGAGCAGGG (reverse). The *Mbd2* mice were genotyped using a combination of three primers: TTGTGAGCTGTTGGCATTGT, GTCAACAGCATTTCCCAGGT and TGTCCTCCAGTCTCCTCCAC. The wildtype *Mbd2* mice showed a single band with a size of 377 bp, the homozygous KOs amplified a single 250 bp product while the heterozygous showed bands for both 377 and 250 bp products when the PCR amplified products were run on an agarose gel.

Once the animals start to develop palpable mammary tumors spontaneously, the size of the tumors were measured at weekly intervals using a Vernier caliper and the tumor volume was calculated by the following formula: V= (length × Width^2^)/2, as described by us before (48).

### RNA extraction and quantitative polymerase chain reaction (qPCR)

RNA extraction was done from frozen tissues. Total RNA was extracted using the AllPrep DNA/RNA Mini Kit (Cat# 80204, Qiagen, Hilden, Germany). The RNA concentration was measured using BioDrop, and 2 μg of total RNA from each sample was used for reverse transcription-polymerase chain reaction (RT-PCR) with random hexamer primers (Invitrogen, Waltham, MA, USA; Cat#58875). SYBR^®^ Green (Applied Biosystems, Cat#A25742) based quantitative PCR (qPCR) assay was performed using an ABI StepOnePlus™ (Applied Biosystems) machine. The changes in gene expression between the control and different treatment groups were determined using the 2^-ΔΔC^T method.

### RNA-Seq and analysis pipeline

RNA extracted from the mammary tumors of PyMT; *Mbd2^+/+^* and PyMT;*Mbd2^-/-^* animals were subjected to RNA-Sequencing (n=3/group). First, the integrity of the RNA was checked and confirmed with an Agilent 2100 Bioanalyzer, and then ribosomal RNA was removed from the samples by the Ribo-Zero kit (Illumina, San Diego, CA, United States). The sequencing library was prepared following the standard protocol for TruSeq Stranded Total RNA Sample Prep Kit (Illumina), and paired-ended [2×150 bp] sequencing was performed on NovaSeq 6000 sequencing system (Illumina). Once the sequencing run was completed, the adaptor sequences, low quality and undetermined bases were removed, and the quality of the reads was verified by FastQC (http://www.bioinformatics.babraham.ac.uk/projects/fastqc/). The de-multiplexed reads were then mapped to the reference genome of *Mus musculus* (Version: v90) using Bowtie2 (49) and HISAT2 (50) aligners. Assembly of the mapped sequencing reads and differential expression of the transcripts were estimated using StringTie (51) and edgeR (52). The known long non-coding RNAs (lncRNA) were identified based on sequence similarities. To identify the novel lncRNAs, we first filtered out the transcripts having an overlap with known mRNAs, known lncRNAs, and transcripts shorter than 200 bp. Then Coding Potential Calculator (CPC) (53) and Coding-Non-Coding-Index (CNCI) (54) tools were used for predicting the transcripts having coding potential. The transcripts having CPC score <-1 and CNCI score <0 was then filtered out, and the remaining transcripts were considered as novel lncRNAs. The known and novel lncRNAs were then combined together and used for downstream analyses. For both mRNAs and lncRNAs, the transcripts passing the following two criteria: (1) log_2_ (fold change) greater than 1 or log_2_ (fold change) less than −1, and (2) *P*-value < 0.05 (parametric F-test comparing nested linear models) were considered as differentially expressed. Pathway analyses were performed by using ConsensusPathDB (55).

### Immunoblotting

Immunoblotting was performed from snap-frozen mouse primary tumor tissues and cell lines. Like the RNA extraction procedure, frozen tumors’ processing had an extra homogenization step under cryogenic conditions. We used a radioimmunoprecipitation assay (RIPA) buffer supplemented with a mixture of appropriate protease and phosphatase inhibitors to prepare cell lysates. Following quantification, an equal amount of proteins was electrophoresed on 8 to 15% sodium dodecyl sulfate-polyacrylamide gels prepared in-house and transferred to polyvinylidene difluoride (PVDF) (Bio-Rad, Hercules, CA, USA; Cat# 1620177) membrane at 4°C. The membrane was soaked into 5% milk in Tris-buffered saline (TBS) to block non-specific antibody binding. Then appropriate primary and secondary antibodies were used, and enhanced chemiluminescence detection kit (Amersham, GE Healthcare Life Sciences, Cat# RPN2232) was used for the visualization of different proteins by a ChemiDoc MP Imaging System (Bio-Rad Laboratories, Inc., Hercules, CA, United States).

### Proteomics analyses of the tumor samples

Cell lysates obtained from the homogenized mammary tumors of PyMT; *Mbd2^+/+^* and PyMT;*Mbd2^-/-^* animals were subjected to proteomic profiling using an Ultra performance liquid chromatography-tandem mass spectrometer (uHPLC/MS-MS) at the RI-MUHC proteomics core (n=3/group). For the identification of peptides and proteins, Scaffold (version 4.9) was used (56). The cut-off probability for peptide identification was set at 90% minimum. For the identification of proteins, the threshold was set at 95.0% minimum and at least two peptides minimum. For identification of protein with differential abundance in the two groups, a *P*-value cut-off of less than or equal to 0.05 was considered statistically significant.

### Ki67 staining

The formalin-fixed mammary tumor tissues were stained with a monoclonal antibody against Ki67 (Dako, Cat# M7240), and the number of ki67 positive cells was determined from the photomicrographs of five randomly selected fields for each sample by ‘ImmunoRatio’ (57).

### Statistical analyses

Results in different graphical representations are shown as the mean ± standard error of the mean (SEM) unless mentioned otherwise. Depending on the number of groups during analyses, a student’s *t*-test or ANOVA was done to measure statistical significance. A *P*-value of less than or equal to 0.05 was considered statistical significance.

## Acknowledgement

This study was supported by grants from the Canadian Institutes for Health Research ‘MOP 130410’ to SAR. and MS and ‘PJT-156225’ to SAR. NM is the recipient of Fonds de la recherche en santé du Québec (FRQS) Doctoral fellowship from the Government of Quebec, Canada.

