## Supplementary Figure for "Methyl-CpG binding domain protein 2 plays a causal role in breast cancer growth and metastasis"

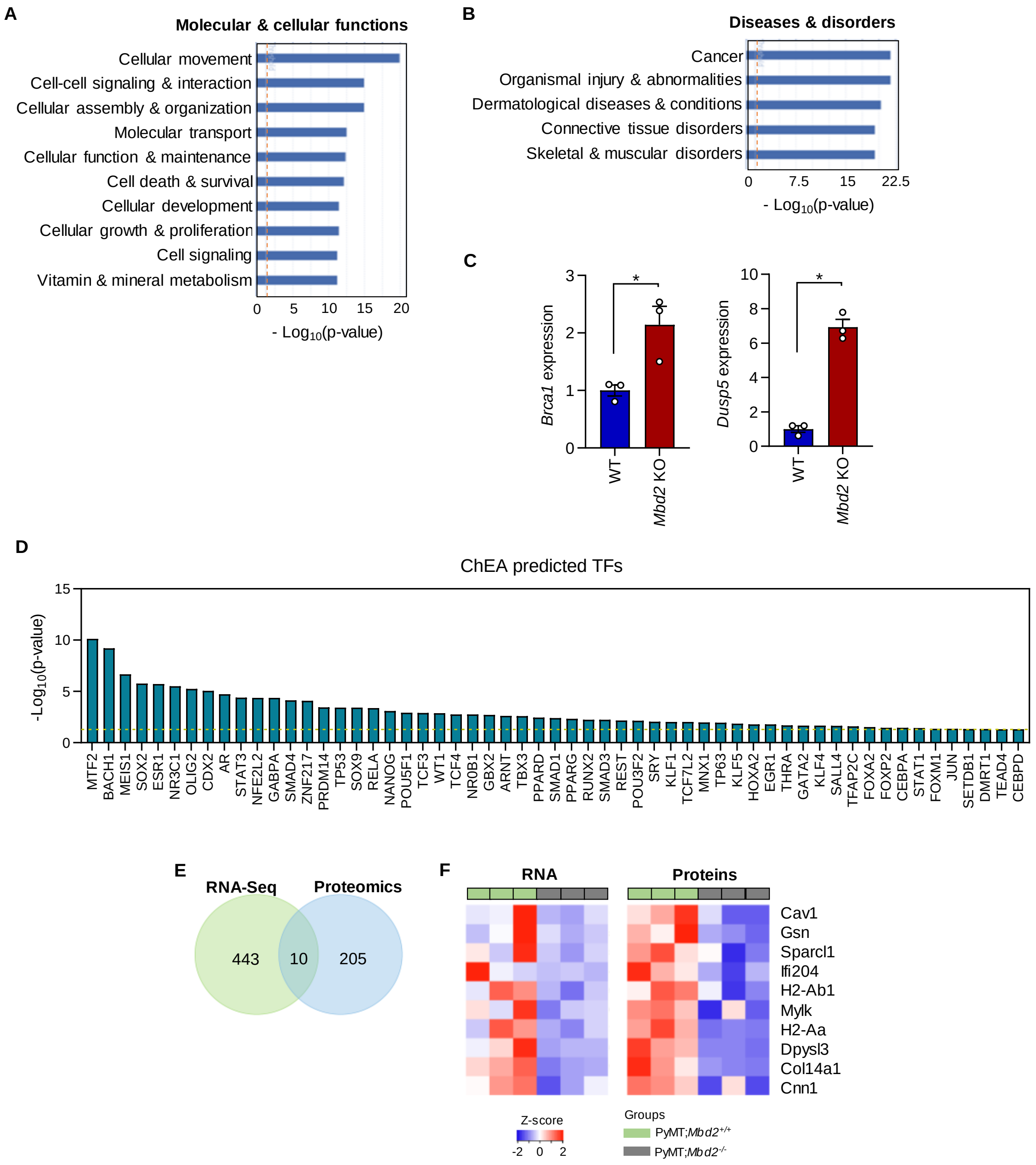


**Supplementary Figure S1: Transcriptomic and proteomic analyses. (A)** IPA tool generated top ten 'molecular and cellular functions' that are altered by the DEGs in *Mbd2* KO tumors. **(B)** IPA generated top five 'diseases and disorders' altered by the DEGs in *Mbd2* KO tumors. **(C)** qPCR validation of the selected tumor suppressor genes (*Brca1, Dusp5*) obtained from RNA-Seq was performed using tumoral RNA from at least three animals/group. **(D)** ChEA predicted upstream transcription factors for the DEGs identified in *Mbd2* KO tumors. **(E)** Venn diagram showing the overlap of 10 genes between RNA-Seq and proteomics analyses. **(F)** Heatmap showing the overlapped genes showed concordant downregulation in their expression in both platforms.

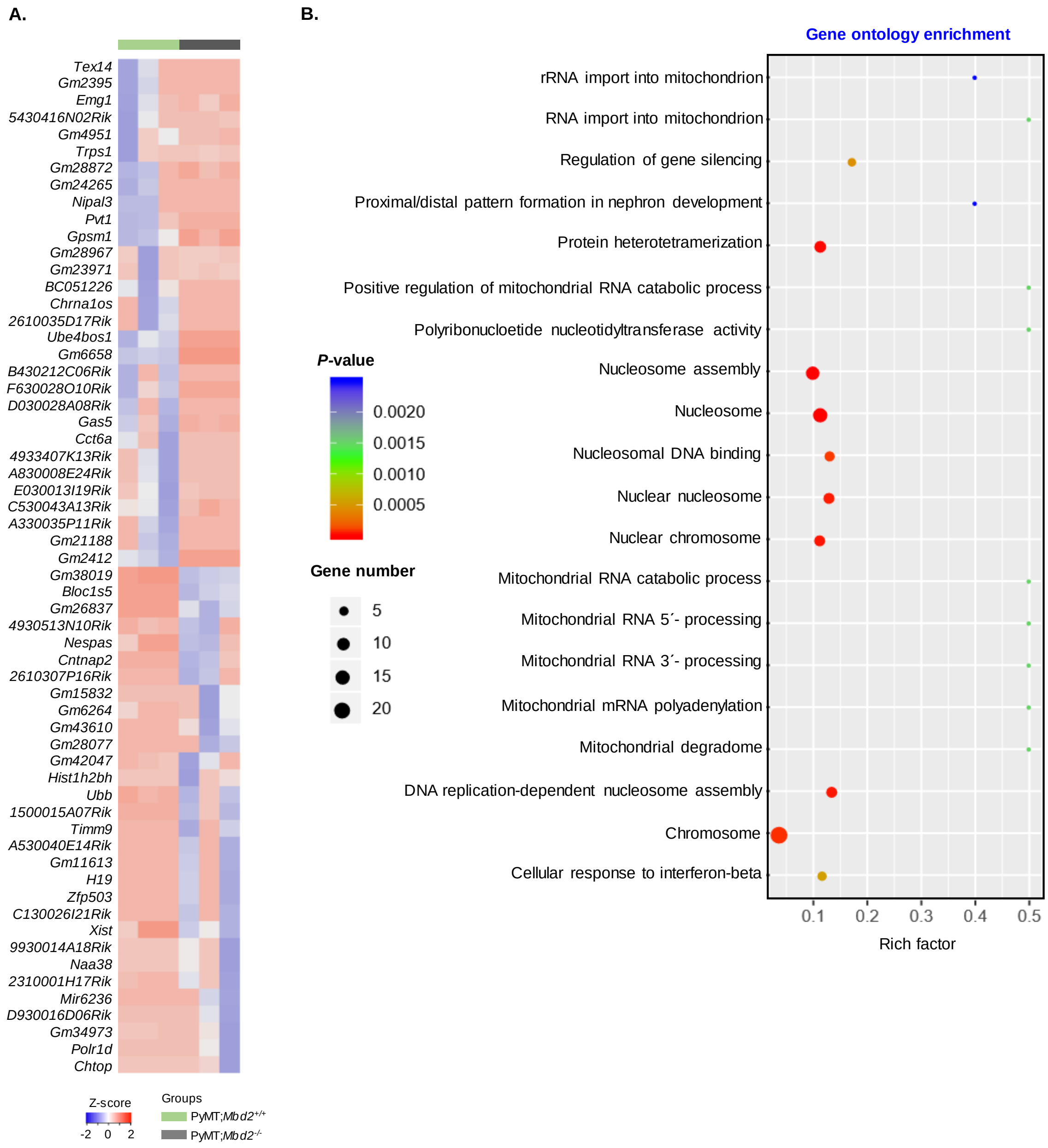


**Supplementary Figure S2: *Mbd2* regulated lncRNAs. (A)** Heatmap of the 60 differentially regulated lncRNAs in Mbd2 KO tumors. **(B)** Gene ontology (GO) enrichment analyses using the list of differentially regulated lncRNAs.
